# Expression profile analysis of circular RNAs in essential hypertension by microarray and bioinformatics

**DOI:** 10.1101/376178

**Authors:** Xingjie-Bao, Xin-He, Shuying-Zheng, Yizhe-Luo, Tianlun-Gu, Ronghui-Tan, Jihan-Sun, Jinshun-Zhao, Lina-Zhang

**Affiliations:** Department of Preventative Medicine, Zhejiang Provincial Key Laboratory of Pathological and Physiological Technology, Medicine School of Ningbo University, 818 Fenghua Road, Ningbo, Zhejiang Province, 315211, China

**Author notes:** Corresponding author E-mail address (Lina Zhang).

**Keywords:** essential hypertension, bioinformatics, circular RNAs, microarray, network

## Abstract

Circular RNAs (circRNAs), widely found in human cells, are involved in disease and play an important role in progression. To determine whether circRNAs are related in essential hypertension (EH), we analyzed the expression profile of circRNAs and miRNAs in 5 EH and 5 healthy controls cases which were screened by microarray. Through microarray data and public data analysis, differently expressed transcripts were divided into modules, and circRNAs were functionally annotated by miRNAs. The expression of two circRNAs, has_circ_0037909 and has_circ_0105015, were validated in EH by qRT-PCR, which may be associated with EH. Further analysis showed that two circRNAs might through immune system by up-regulation circRNAs and down-regulation expression. These circRNAs biological functions need to be further validated.

## 1 Introduction

Essential hypertension (EH) is a common disease that threatens the health of the population, characterize by an abnormal increase systolic blood pressure (SBP) and/or diastolic blood pressure (DBP). With the development of molecular diagnostic technology, more accurate prediction and diagnostic techniques have been applied to the clinic and brought a gospel to patients. The rapid evolution of high throughput technologies (HTS) including cDNA microarray and next-generation sequencing has founded some new mutations, such as CYP2C19 (Choi *et al.* 2014) and ERAP1 (Yang *et al.* 2015) in EH. However, most of transcripts detected by HTS were not protein-coding genes but non-coding RNAs. And there are no protein-coding mutations in some diseases, but instead of non-coding RNA e.g. small non-coding RNAs. Circular RNAs (circRNAs) may be the main character.

We know about circRNAs by using it as sponges of microRNA (miRNAs), adjust splicing and transcription, and gene expression modify (Qu *et al.* 2015). Studies have shown that circRNAs may play an important role in cardiovascular disease(Fan *et al.* 2017). In adult murine hearts, circRNAs are key roles of bioinformatics and sequencing technology (Jakobi *et al.* 2016). And they are highly specific to cell tissue and developmental status(Salzman *et al.* 2013). The effect of a heart-related circRNA on the protection of the heart from heart failure and pathological hypertrophy was targeted by miR-223 and as sponges of miR-223(Wang *et al.* 2016). Even recently our team research shows that circRNAs may have an effect on the hypertension. Has_circ_0037911 is a potential risk factor of EH via changing the amount of Serum creatinine in the body (Bao *et al.* 2018). It is expected that circRNAs will be performed for clinical diagnosis and accurate prediction in large-scale clinical trials. our study analyzed the expression profiles of circRNAs and miRNAs in EH and health control, and focused on the important role of circRNAs in EH in the process of modularization. And we identified the functions of circRNAs by using public databases. It is likely that methods of microarray and bioinformatics of the genome will improve our understanding of the contribution in EH.

## 2 MATERIALS AND METHODS

### 2.1 Patients and samples

In the first step, the whole blood samples were obtained from 5 cases with essential hypertension and 5 control group cases at seventh people’s hospital in 2016. All research subjects informed and understanding the written consent before enrolled into the study. Diagnosis of hypertension was made in accordance with the 2016 guidelines and management of hypertension in adults (Gabb *et al.* 2016): systolic blood pressure (SBP) > 140mmHg and/or diastolic blood pressure (DBP) > 90 mmHg; controls with SBP < 120mmHg and DBP < 80mmHg. All cases had no history of secondary hypertension, diabetes mellitus, myocardial infarction, stroke, renal failure, drug abuse or other serious diseases, no family history of hypertension in the first degree relatives, and did not receive treatment for hypertension. The baseline characteristics of the 10 samples are shown in table 1. In the second step, we obtained 20 EH cases and 20 sex- and age- (±3 years) match healthy controls, and validate performed the real-time quantitative reverse transcription-polymerase chain reaction (qRT-PCR). After 12 hours of fasting, blood samples were collected from the antecubital vein using ethylenediaminetetraacetic acid (EDTA) and stored at -80 °C.

**Table 1.** The baseline characteristics of the 10 samples. M, male; F, female.

### 2.2 Profiling of circRNA expression

The expression of circRNAs was profiled with CapitalBio Technology Human CircRNA Array v2 conducted by CapitalBio technology (Beijing, China). Total RNA samples from peripheral blood were transcribed into first strand cDNA and second strand cDNA. Then, the CapitalBio Technology Human CircRNA Array v2 were hybridized each cDNA. Designed for the global expression profiling of human circRNAs and miRNAs transcripts, this array can detect circRNAs in two hierarchically compiled: standard circRNAs, used for functionally-enhanced functional studies and experimental support of complete circRNAs, and for reliable high a comprehensive collection of circRNAs for confidence. The circRNAs were carefully constructed using landmark publications and the most recognized public transcriptome databases (eg, circBase, known genes and collections of miRBase).

### 2.3 Mapping and identification of differentially expressed genes

We employed Agilent Feature Extraction (v10.7) software to analyze hybrid images and extract data. Then applied Agilent GeneSpring software to normalize and analyze the data: fold change >2 for up-regulation or down-regulation; *p*-value <0.05; and false discovery rate (FDR) < 0.05.

### 2.4 Gene Ontology and Kyoto Encyclopedia of Genes and Genomes pathway analyses

Gene Ontology (GO) analysis was intended to facilitate the consistent description of the function of gene products in various databases. According to the latest Kyoto Gene and Genome Encyclopedia (KEGG) database, pathway analysis of differently expressed genes was studied to find important pathways. The significant pathways and GO terms were determined by Fisher’s exact test, and FDR was used to correct the *p*-values.

### 2.5 Validation of circRNAs by qRT-PCR

Expression of five circRNAs verified carried out by qRT-PCR. The synthesized of circRNAs-cDNA was performed a GoScriptTM Reverse Transcription System kit (Promega, Beijing, China), the miRNAs-cDNA was miRcute Plus miRNA First-Strand cDNA Synthesis Kit (TianGen, Beijing, China). QRT-PCR was conducted with the GoTaq^®^ Real-Time PCR Systems reagent kit (Promega, Beijing, China), and miRcute Plus miRNA Real-Time PCR Systems reagent kit (TianGen, Beijing, China) following the manufacturer’s instructions. All primers were synthesized by Invitrogen (Shanghai, China).

### 2.6 Statistical analysis

The expression of circRNAs was analyzed by the ΔCt method using SPSS software 19.0(Yang *et al.* 2016) (SPSS Inc., Chicago, IL, USA). Differences of circRNAs levels between EH group and control group were assessed using t-tests. A two-side *p* < 0.05 was considered statistically significant. The ROC curve was use to show the diagnostic value more deeply.

## 3 RESULTS

### 3.1 Differentially expressed circRNAs and miRNAs in AML

Volcano plots were used to assess gene expression variation between case and control groups. A total of 287 circRNAs showed differential expression in both groups, including 287 up-regulated circRNAs and 78 down-regulated circRNAs. The top 20 circRNAs were significantly up-regulated or down-regulated among two groups. Hierarchical cluster analysis also showed systematic changes in circRNAs expression. (Figure 1).

**Fig 1.**
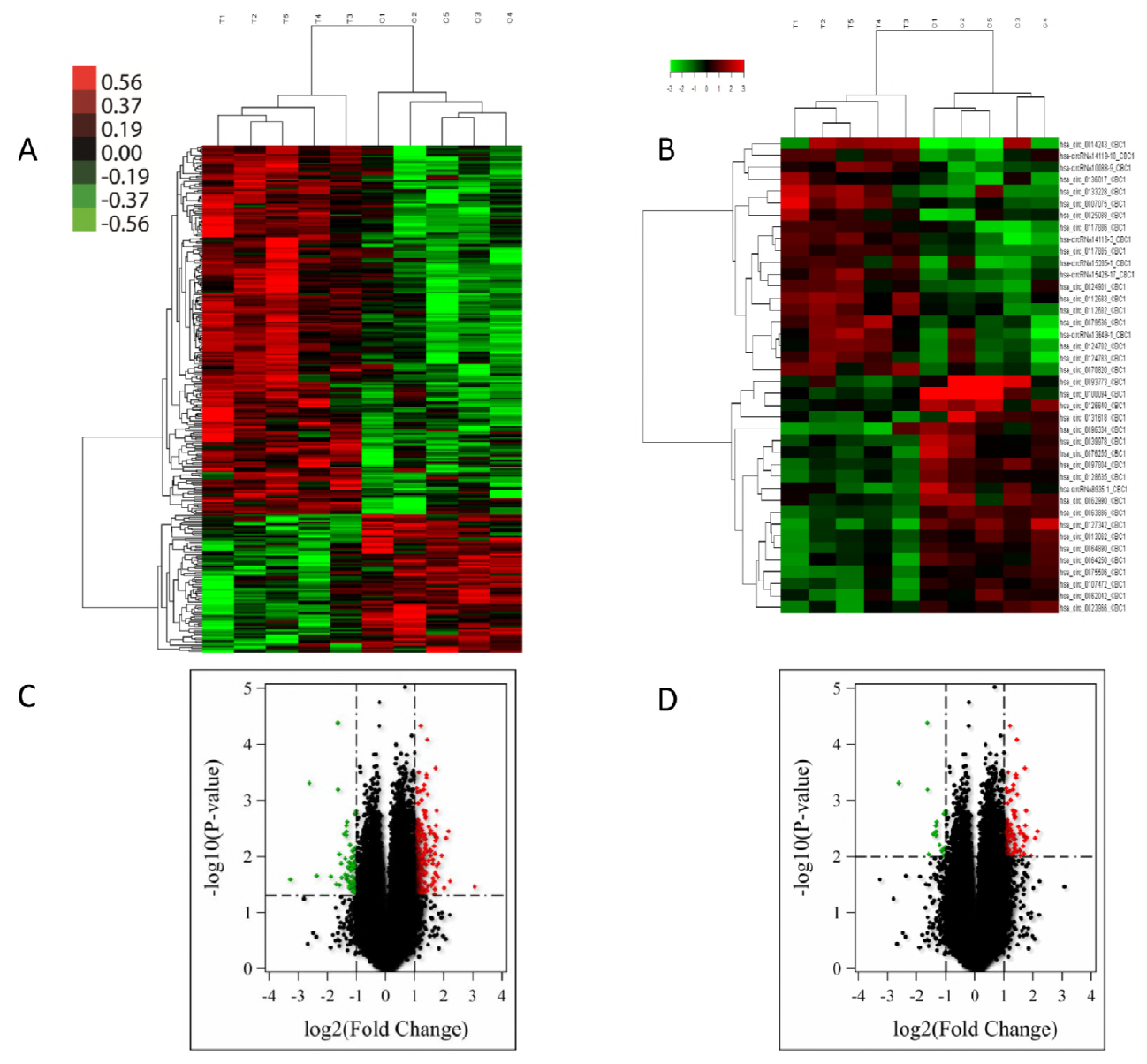
Volcano plots and heat map showing expression profiles of circRNAs in essential hypertension. A and B, maps showing the hierarchical clustering and differential expression of circRNAs between 5 paired samples; C and D, panels showing significantly changed circRNAs with p-value < 0.05 and p-value <0.01.

These circRNAs are widely distributed in 24 chromosomes (Figure 2A). The total circRNAs were divided into two categories: 287 up-regulated circRNAs and 78 down-regulated circRNAs. The top 10 of circRNA-miRNA-network were show in figure 2B. Top 20 up-regulated and down-regulated circRNAs in hypertensive patients were be showing in table 2 and table 3.

**Fig 2.**
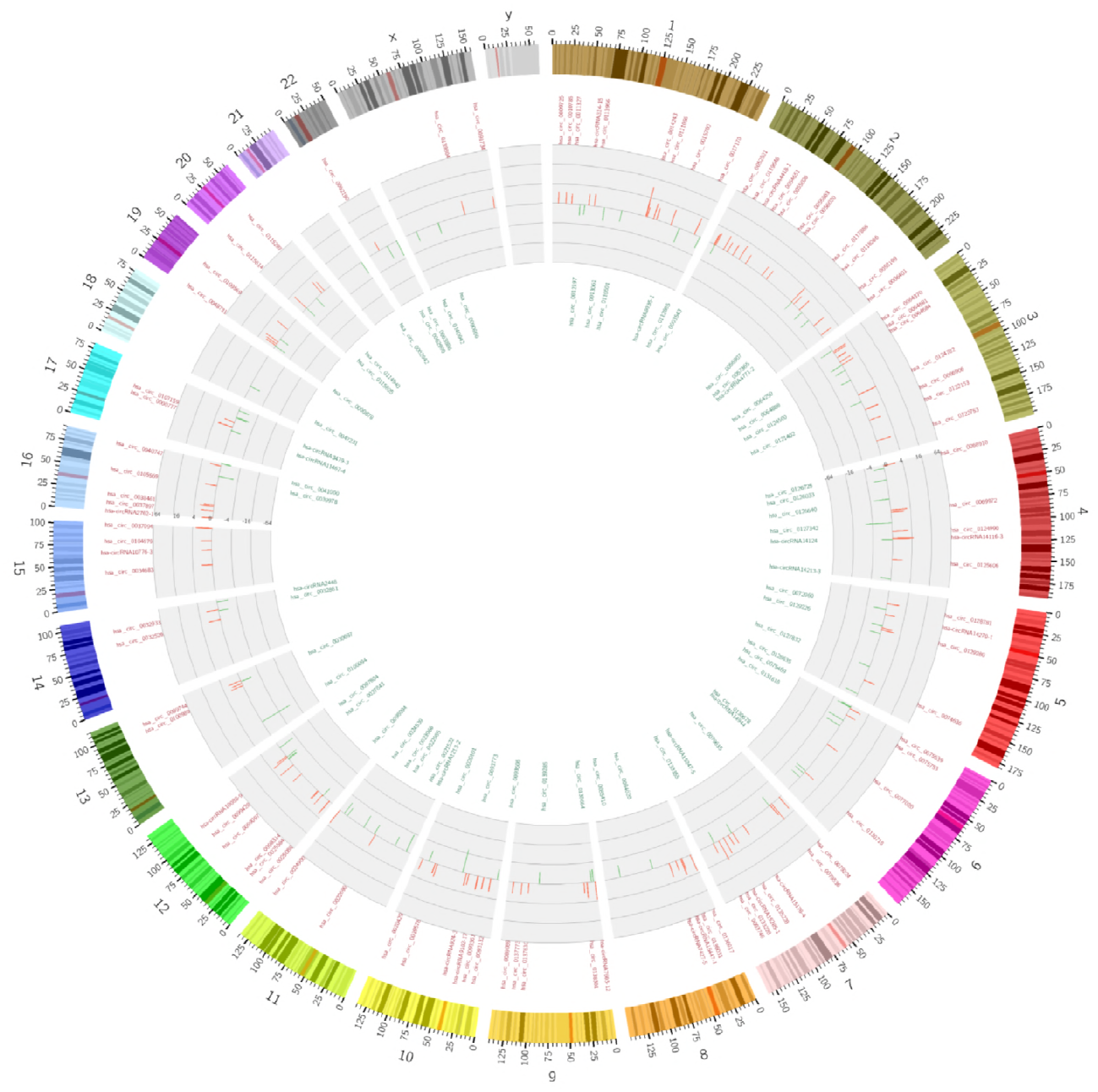
Identification of differentially expressed circular RNAs (circRNAs) in essential hypertension (EH). A, Circos plot showing circRNAs on human chromosomes. From the outside in, the first layer of the Circos plot is a chromosome map of the human genome, black and white bars are chromosome cytobands, and red bars represent centromeres. In the second circle, all up-regulated circRNAs were showed and all down-regulated circRNAs were in the third layer; B, the top 10 of circRNA-miRNA network maps the interaction relationship between circRNAs and miRNAs.

**Table 2.** Top 20 up-regulated circRNAs in hypertensive patients.

**Table 3.** Top 20 down-regulated circRNAs in hypertensive patients.

**Table 4.** Comparison of Characteristics between 48 controls and 48 EH group.

### 3.2 Functional analysis of differentially expressed genes

There have not yet been well annotated the function of most circRNAs. Therefore we can predict the role of circRNAs in EH by GO and KEGG pathway analyzing, and be provided with a clue about the EH disease process. According to Go-Standard, the circRNAs were divided into three parts of functions including biological process, celluar component and molecular function (Figure 3A). In the KEGG pathway analysis, the significant enriched of expression of circRNAs were found in ECM proteoglycans and Extracellular matrix organization pathway terms (Figure 3B). The cyclic-nucleotide phosphodiesterase activity and hemostasis pathway terms were found in GO pathway analysis (Figure 3C).

**Fig 3.**
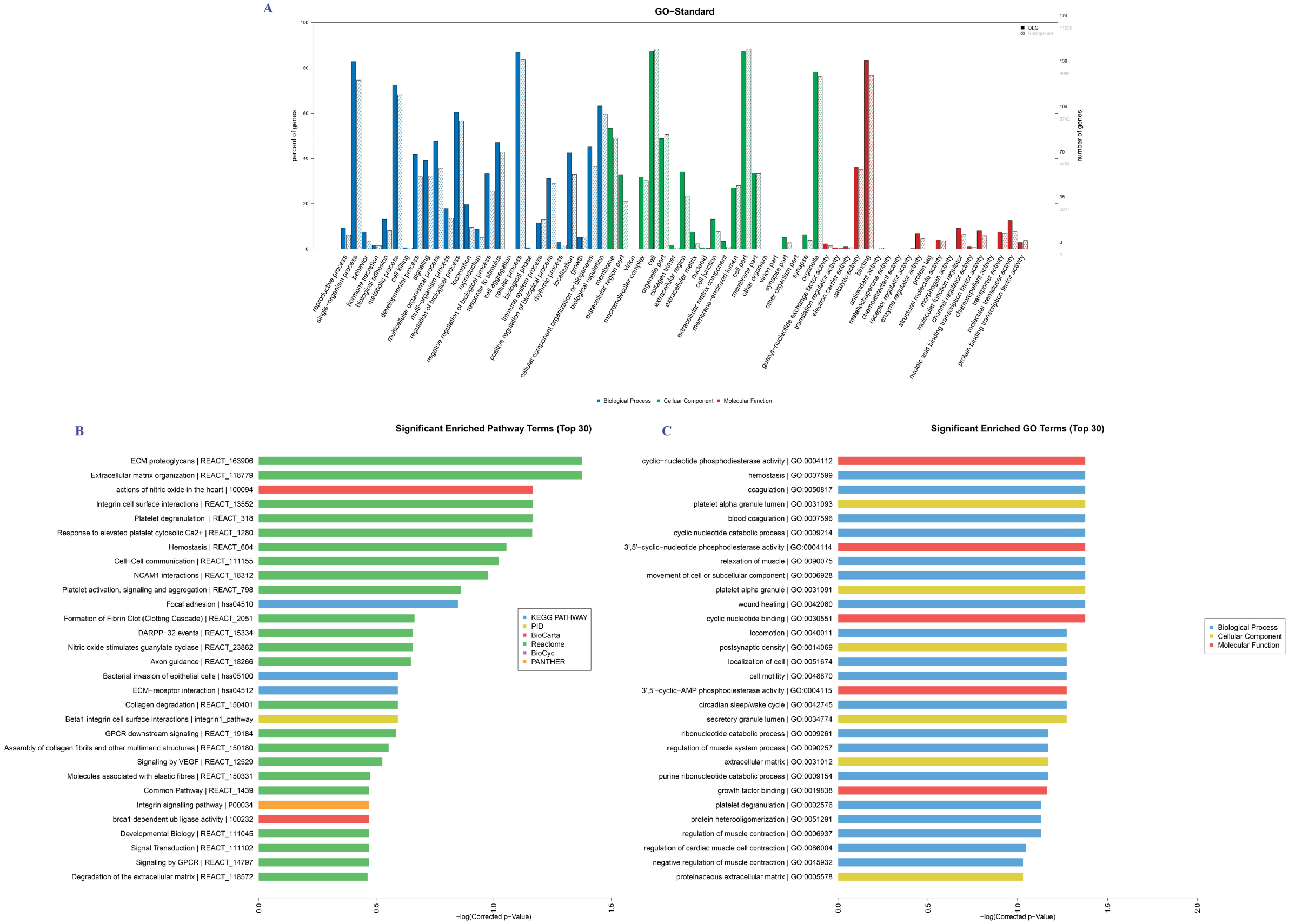
Gene Ontology (GO) and Kyoto Encyclopedia of Genes and Genomes (KEGG) pathway analysis of circRNAs in essential hypertension. A, all circRNAs were divided into three part of function: biological process, celluar component and molecular function; B, KEGG pathway analysis about top 30 of significant enriched pathway terms; C, GO pathway analysis about top 30 of significant enriched GO terms.

**Fig 4.**
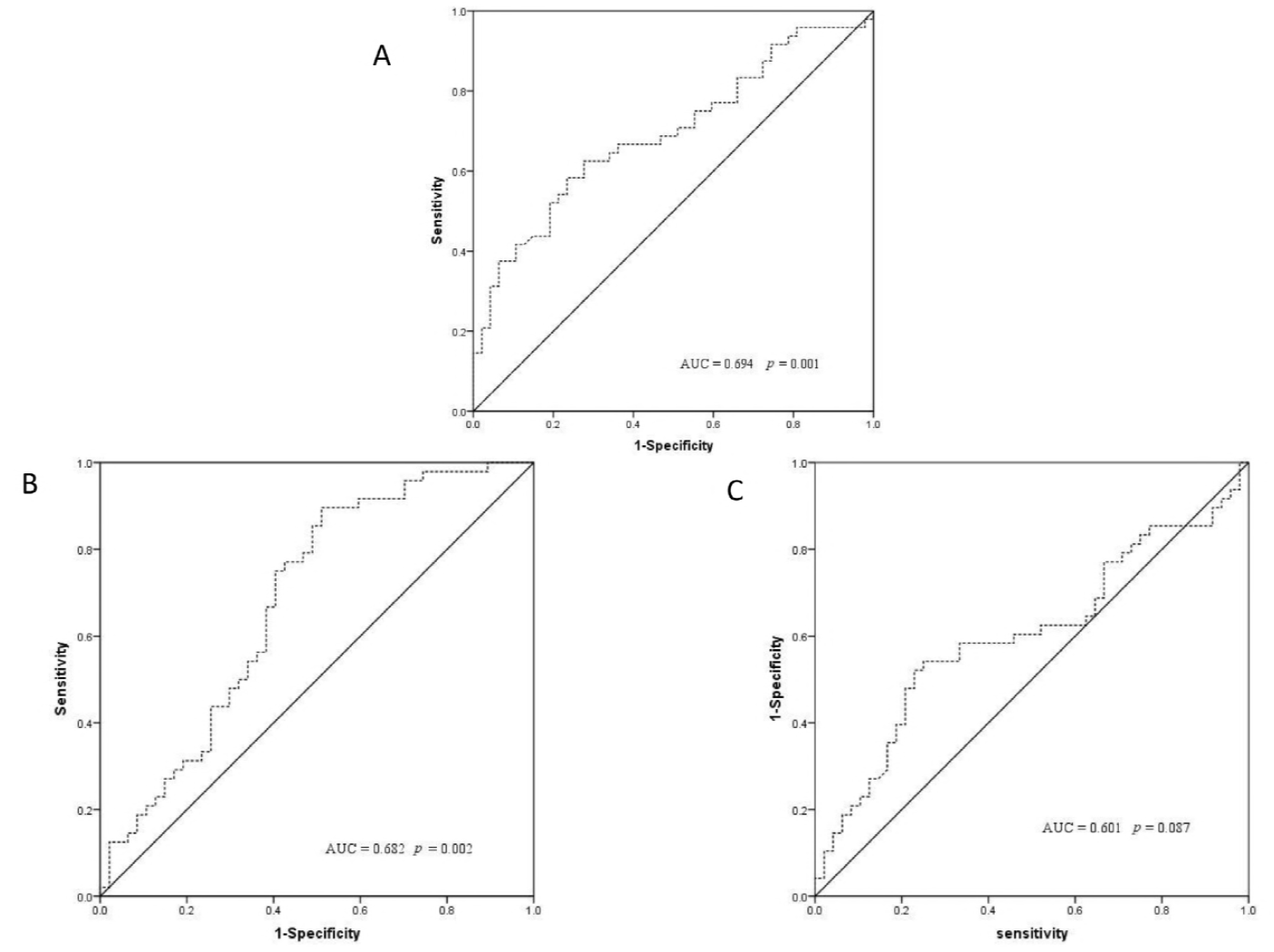
The ROC curve for circRNAs and miRNA. A, the ROC analysis for hsa_circ_0105015; B, the ROC analysis for hsa_circ_0037909; C, the ROC analysis for has-miR-637.

### 3.3 Validation of regulated circRNAs in EH and healthy controls

We used qRT-PCR to detect circRNAs expression in order to validate microarray data. Hsa_circ_0105015 and hsa_circ_0036909 are up-regulated in microarray data and has-miR-637 is down-regulated. Through our validate phase, the same results as microarray data were confirmed in 48 EH and 48 healthy control cases. Furthermore, we can conclude that circRNA can act as sponges of microRNA and reduce its expression.

### 3.4 Sensitivity analysis using receiver operating characteristics (ROC) curve for circRNAs

After the systemic analysis of circRNAs and miRNA expression profiling, combine GO and KEGG data which provides genetic studies of specific disease associations and extensive human gene expression, we have confirmed the core circRNAs and found some circRNAs may involve the occurrence of EH patients. In order to determine whether circRNAs have diagnostic value, the ROC analysis was carried out. The results showed hsa_circ_0105015 and hsa_circ_0037909 were significant to predict diagnostic. The AUC were 0.694(*p* = 0.001) and 0.682 (*p* = 0.002), respectively. In summary, these two circRNAs not only have abnormally expression in EH, but also had important clinical significance. Further functional studies of these circRNAs would be of value.

## DISCUSSION

It is well known that multiple factors (e.g. genes, behaviors, surrounding) are the characteristics of EH (Karaca *et al.* 2014). As new biological roles, circRNAs have only just been understood, while most of them have not been studied yet. We conducted a comprehensive analysis of the circRNAs and miRNAs profiling data of patients with EH and healthy controls, and performed a circRNA-miRNA-network model together with a public database, in order to thoroughly exchange the biological work of circRNA in the pathogenesis of EH. In short, circRNAs play a very important role in our established circRNA-miRNA-network model among EH.

We investigated the expression profile of transcripts between peripheral blood of EH cases and healthy controls. We used microarray method to identify large numbers of circRNAs and miRNAs, supporting widespread participation in circRNAs in EH. The mass of data was handled in two phases with two different important concepts. We have simplified the composite transcription network through modules in first phase. MiRNAs were divided into several functional modules indicating the effectiveness of microarray by GO and KEGG pathway, including cardiovascular diseases, arrhythmogenic right ventricular cardiomyopathy (ARVC), congenital muscular dystrophies (CMD/MDC), hypercholesterolemia, fanconi anemia and long QT syndrome. In addition, the pathway is divided into focal adhesion, bacterial invasion of epithelial cells and extracellular matrix (ECM) receptor interaction in significant enriched pathway terms. In conclusion, the above method is helpful to deeper understand “big data”. In second phase, the dataset from circBase and miRBase was conducted to analysis the clinical significance of circRNAs from the modules. Two circRNAs, including has_circ_0037909 and has_circ_0105015, may serve as biomarkers for EH and predicting the occurrence of disease. According to their correspondingly adsorbed miRNAs and belonging to the most important network, the expression of has_circ_0037909 and has_circ_0105015 are related to the operating system of EH. Combining public data and some studies (Nemecz *et al.* 2016), circRNAs serve as cavernous bodies of miRNAs and may regulate inflammatory pathways by affecting the expression of miRNAs to damage the endothelial cells.

Our work clearly reveals that circRNAs are an important role in EH. Most of the circRNAs, however, were not assigned to any functional module and were not included in public data and scientists’ research. In the initial stages of study, the functions and targets of circRNAs which may also play a crucial role in EH were difficult to understand. In addition, the results also show than circRNA-adsorbed miR-637 was associated with the vascular immune system including C-reactive protein (Kim *et al.* 2015). CircRNAs may affect the corresponding function by regulation the abundance of the miRNA attached to the body, i.e., as a miRNA aponge. The biological function of circRNAs needs further verification. With the development of research on the biological function of non-coding RNA, we have formed a kind of understanding that RNA is not a simple garbage link, but directly involved in the regulation of biological networks (Conn *et al.* 2015; Bonizzato *et al.* 2016). With the development of research techniques – the next generation of sequencing in terms of depth and scale, we have accumulated a lot of data. Through the above technologies, we realize that this system is complex and exceeds the scope of initial recognition range. The good news is advances in methods have simplified the network. By modularizing and simplifying the network, a large number of transcripts are converted into several great pathways, and then the core content is studied according to each pathway. Moreover, we will actively apply the accumulated public data, which may help us to study the function of circRNAs. And we will identify functional annotations of core circRNAs and employed qRT-PCR to verify their expression.

## ACKNOWLEDGMENTS

This work was supported by the National Natural Science Foundation of China (Grant No. 81773528); Ningbo Scientific Innovation Team for Environmental Hazardous Factor Control and Prevention (Grant No. 2016C51001); Zhejiang Province Social Development Research Project (Grant No. 2016C33178); K.C. Wong Magna Fund in Ningbo University, Ningbo Social Development Research Project (Grant No. 2014C50051).

## CONFLICT OF INTEREST

The authors have no conflict of interest.

